# Estimating the completeness of large-scale single-cell sequencing projects

**DOI:** 10.1101/2025.01.08.631769

**Authors:** Mitro Miihkinen, Yidian Chu, Sara Vakkilainen, Yevhen Akimov, Tero Aittokallio

**Author notes:** These authors contributed equally to this work (Yidian Chu, Sara Vakkilainen).

## Abstract

During embryonic development, cells undergo differentiation into highly specialized cell types. Capitalizing on single-cell RNA sequencing, many initiatives and substantial resources have been established for cataloguing these differentiated cell types by their transcriptomic profiles. Despite the extensive efforts to profile various organs and their cellular compositions, we lack metrics to assess the completeness of the sequencing projects. In this cellular biodiversity analysis, we leveraged the increasingly available single-cell data together with statistical methods, originally developed for assessing the species richness of ecological communities, to estimate the cellular diversity of any organ based on current data from single-cell profiling technologies. Deriving from such cellular richness estimates, we established a practical statistical framework that enables reliable assessment of the completeness of any large-scale single-cell profiling projects, after which additional sequencing efforts do not anymore reveal new insights into an organ’s cellular composition. Such estimates can serve as stoppage-points for the ongoing sequencing projects, hence guiding a more cost-efficient completion of the profiling of various human tissues.

## Introduction

Each organ in a human body can be thought of as a natural community, an ecosystem, composed of various, morphologically and functionally diverse cell types. In an adult human, differentiated cells are defined by their cell type-specific spatial location, function and gene expression, which both defines and maintains cell identity and function (Zeng, 2022). For this reason, the emergence of single-cell RNA sequencing has paved the way for profiling and cataloguing cells from different organs, placing them into increasingly granular cell type categories. Even though there does not exist a clear consensus on the level of granularity at which cell types should be classified, such cataloguing efforts and ambitious international initiatives, such as the human cell atlas, are aiming to build comprehensive cell type reference maps as a basis for understanding human health and disease (Rood et al., 2024). However, the lack of accurate estimates of the number of unique cell types make it challenging to know how complete such cataloguing efforts currently are.

The cellular communities that compose different organs can be analysed using biodiversity metrics. Species richness of an ecosystem, or any natural community, is the simplest and most intuitive description of biodiversity, which is often related to ecosystem function and resilience (Gwinn et al., 2016). However, identifying and cataloguing every individual in a population to understand species richness is often impractical, expensive and unnecessary (Magurran, 1988). A more effective approach to study the number of species is through replicate sampling of the ecosystem. Since the number of species present in a sample is almost always an underestimate of the true population value, many statistical methods have been proposed to correct this bias and provide more accurate estimates of the species richness of an ecological population (Gwinn et al., 2016). We argue that such estimates could be used also to assess the richness of cellular ecosystems, and to provide completeness estimates for the single-cell atlases, provided the statistical methods are applicable to cellular systems.

Methods to infer species richness from a sample of an ecosystem range from purely data-analytical approaches to parametric and non-parametric statistical models, along with upsampling methods (Schmitz & Rahmann, 2024). While these approaches have been extensively applied to ecological populations—such as animal or plant species—special considerations are required when selecting appropriate estimators for cellular communities, particularly in the context of single-cell or single-nucleus RNA sequencing data. Importantly, most of the statistical models pose underlying assumptions on the population structure, which in many occasions make them unsuitable for estimating species richness of a cellular community (Chiu, 2023; Schmitz & Rahmann, 2024). In the absence of a clear consensus on the best models for species richness estimates in cellular systems, a careful analysis of the suitability and accuracy of best-performing models is required to avoid biased completeness estimates.

As single-cell resolution data from different human tissues becomes increasingly available, we considered human tissues as natural communities consisting of cells with defined cell types and used statistical methods to describe their biodiversity characteristics, such as number and distribution across tissues. Our systematic analysis reveals many key insights into how cell types are distributed along different organ systems, which together with statistical modelling give an informative means to guide experimental single-cell profiling efforts. We show that cell communities in a human can be generally modelled with a log-linear distribution. Importantly, using single-cell data together with statistical species richness estimation methods, we investigate the completeness of currently available single-cell resources and suggest ways on how statistical analysis can enrich the ongoing efforts to build most comprehensive single cell atlases.

## Results

### Cellular communities in human follow a log-linear distribution

To investigate the cellular biodiversity among different organ systems in a human body, we exploited data of cell type abundances across 20 human organs from Tabula Sapiens (Tabula Sapiens Consortium, 2022), and assessed the statistical properties of each detected cell type across the 20 organs (137 cell types in total, as of June 2025). Such an organ-level analysis demonstrated that there exist various degrees of both common and unique cell types across the organs (Figure 1A-C), and provides a principled means to understand organ development through organ-specific and shared cell types. For example, the large number of different cell types both in the eye and lung highlight the complex functions of these organs (Figure 1A-C), whereas the large number of unique, organ-specific cell types in the eye reflects the evolutionary time needed to develop such a specialized organ (Figure 1A-C). The eye had the most specialized repertoire of unique cell types (58.1% unique cell types), while the lymph nodes had the least unique cell types (4.8%). This was also true for the other organs of the lymphatics system, such as speen (5.9% unique cell types), reflecting the migratory nature of these cells (Figure 1D). Our analysis revealed distinct immune cell populations and fibroblasts as the cell types most common among the organs, reflecting the importance of fibroblasts and the immune system for all organ development and function (Figure 1D). Notably, most cell types (64%) were highly specialized and thus unique to a single organ.

**Fig. 1.**
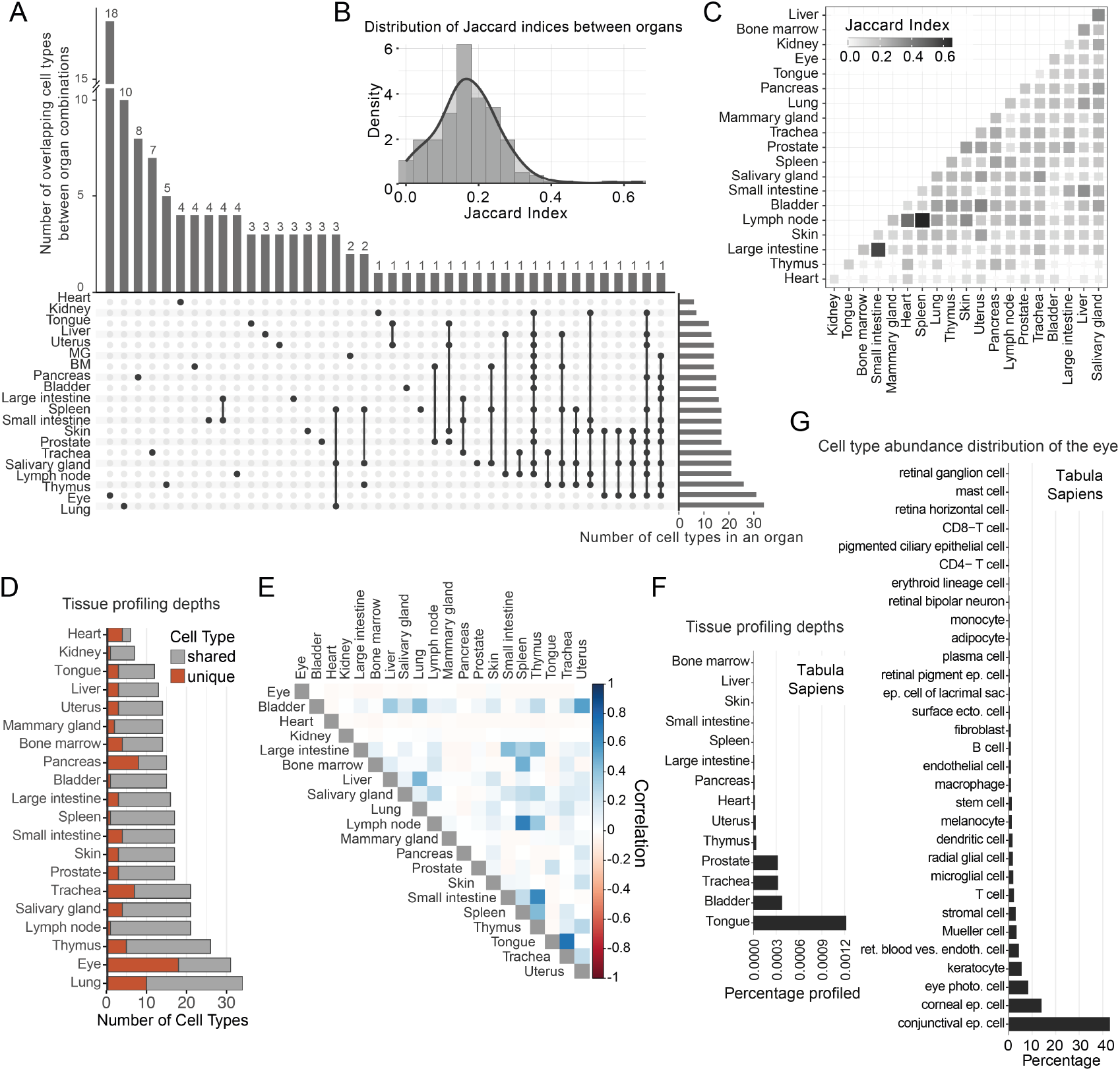
Cellular biodiversity of human organ systems according to Tabula Sapiens data. (A) UpSet plot shows how cell types are distributed across the 20 organ systems in Tabula Sapiens. MG stands for mammary gland, BM stands for Bone Marrow. (B) As a metric for the organ uniqueness, distribution of Jaccard indices are displayed (0 indicates completely dissimilar organs, and 1 completely identical organs). (C) Pairwise Jaccard indices for each pair of organs, computed between each organ’s cell type set. (D) The number of shared and unique cell types across the 20 organs in Tabula Sapiens. (E) Pearson pairwise correlations between organ pairs in Tabula Sapiens dataset, calculated based on the counts of overlapping cell types between the two organs. (F) Depth at which organs in Tabula Sapiens have currently been profiled, calculated as a percentage of the number of cells profiled from the total number of cells in each organ. The estimates of organ total cell numbers are from https://humancelltreemap.mis.mpg.de/ (Figure S1B). Breast tissue is missing since estimates of tissue size were not available (as of June 2024). (G) Lognormal distribution of the eye as an example (KS-statistic 0.07, P=0.99). Test results across all organs shown in Figure S1A.

Since the cell type-specificity analysis demonstrated that cell types can be either shared or unique across organs, we next assessed the similarity of human organs. The organ systems analysed from Tabula Sapiens data showed variable degrees of correlations in terms of the shared cell type compositions (Figure 1E). Although this analysis shows many expected correlations between similar organs (e.g., organs of the lymphatics system, or different parts of the intestine), surprisingly, small intestine and thymus showed unexpectedly high correlation, although not being closely related in terms of their primary function or anatomical location (Figure 1E). Notably, the cell type abundances in the organs displayed a heavy-tailed distribution, and since cell abundances in other cellular ecosystems in bacterial communities have been previously described to follow a log-linear distribution (Grilli, 2020), we tested whether cell type abundances in human organ systems follow this distribution as well. Indeed, Kolmogorov-Smirnov test showed lognormal distribution to be a valid approximation across all the 20-organs tested (Figure S1A). Finally, we wanted to understand the depth at which the individual organs have been profiled. Although sample coverage (the amount of population profiled) is a crucial piece of information to assess the statistical properties of the sample, this is not generally reported in any of the current single-cell atlas projects. We estimated the sample coverage for each of the 20 organs in Tabula Sapiens and observed drastic differences in the levels at which different organs have currently been profiled (Figure 1F). These differences underscore the need for standardized reporting of sample coverage to improve the cross-organ comparisons and highlight the organ systems benefitting from further sampling.

In summary, our analysis of the Tabula Sapiens cell type abundance data gives an overview of the unique and shared distributions of individual cell types for better understanding of these cellular communities. Importantly, we show that organ systems in human are cellular ecosystems whose cell type abundances are approximately log-normally distributed, such as the eye (Figure 1G). In light of previous research done on bacterial species (Grilli, 2020), our analysis supports the log-normal distribution as a natural law from which cellular communities can be derived.

### Statistical framework for estimating cellular species richness

Currently, the completeness of most single-cell atlases is defined simply by the number of cells stored in the database. By such count metrics, the currently largest dataset, CELLxGENE (Program et al., 2023), stores data of 96.1 million cells. Since the human body consists of 35 trillion cells, the completeness of CELLxGENE would be approximately 0.03%. However, since these databases store data from different organs in a non-uniform manner, such estimates are not appropriate for assessing atlas completeness. The count metrics could be improved by displaying them organ-by-organ, but even this approach cannot assess how complete the profiling efforts are. Since one of the immediate goals of single-cell profiling efforts is to exhaustively catalogue all human cell types, we argue that the most relevant way to assess any single-cell atlas completeness is to estimate how many cell types (species) remain to be unrecognized by the current profiling data. Because we lack an established consensus on the definition of cell types, in this work we refer to cell classes as cellular species. To solve this statistical estimation problem (Figure 2A), we assessed the accuracy of different species richness estimation methods when applied to single-cell atlas data using a simulation-based statistical framework (Figure 2B).

**Fig. 2.**
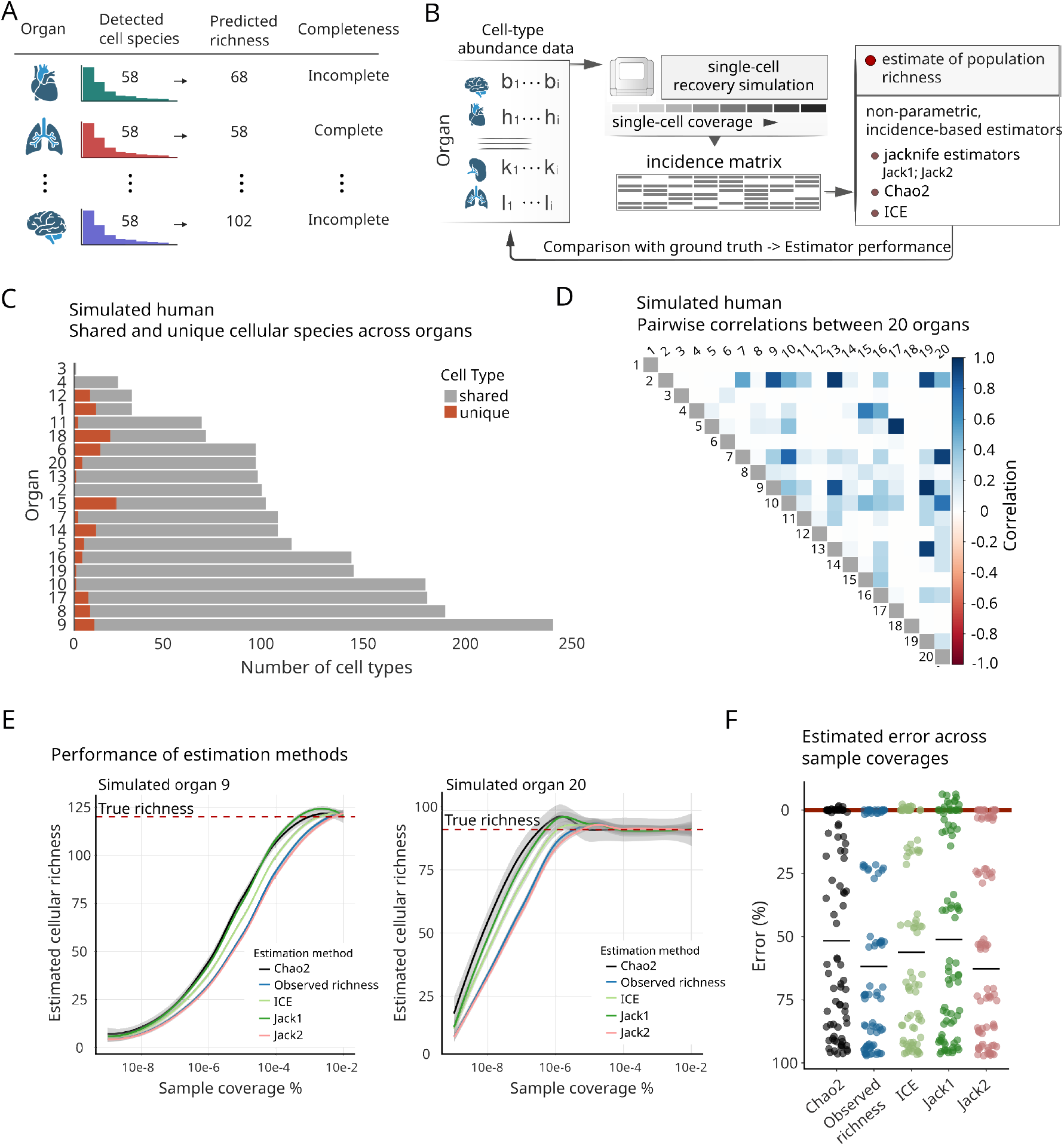
Statistical framework to assess the accuracy of species richness estimators. (A) Statistical framework for any large-scale single-cell project, in which the tissue richness and project completeness are evaluated using species richness statistical estimators. Most existing single-cell atlases and projects report only the total number of cells profiled, offering no insight into undetected cell-type diversity or the overall completeness of the dataset. (B) Schematic illustration of species richness estimator performance testing, where the cell type abundance tables (ground truth population of cells in 20 organs) are established via simulation. The simulation parameters were scaled-up from the Tabula Sapiens data for 20 organs (see Methods for details). For each organ, the abundance vector was repeatedly sampled across six single-cell coverage levels (10^−2^ to 10^−8^) using six bootstrap replicates per level. The resulting incidence matrix was the input for non-parametric richness estimators to evaluate the number of recovered cell types. The estimated richness was compared against the ground truth to calculate percentage of error at each coverage level and LOESS curves. (C) Simulated human: the number of shared and unique cellular species across different organs. (D) Simulated human: Pearson pairwise correlations between different organ pairs, calculated based on the counts of overlapping cellular species. (E) Accuracy of different species richness estimators across different sample sizes from the simulated human. Shown for two simulated organs with different cell species compositions. The true richness indicates the amount of cellular species present in the simulated organ. (F) Error of species richness estimators in a simulated organ profiled at different sample coverages ranging between 0.00000001%-0.01%. The points indicate errors at various sample coverage levels and the horizontal lines the median error levels.

We first established a simulated human body with defined organ systems and cellular species. Since the cell type uniqueness analysis demonstrated that cell types (cellular species) can be either shared or unique across different organs (Figure 1A-C), we simulated realistically distributed and correlated cellular species across all the organs using statistical parameters inferred from the Tabula Sapiens data (see Methods).

We next used the simulated human as the ground truth, from which samples are drawn while accounting for the population complexity and number of cellular species. As expected, the cellular species across simulated organs had distributions similar to those in Tabula Sapiens (Figure 2C). Interestingly, the fraction of cellular species unique to each organ was significantly higher than in the Tabula Sapiens data, providing evidence that many organ-specific species are yet to be discovered (Figure 2C). This is likely true especially for organs which are known to be highly specialized, such as the kidney, which in Tabula Sapiens currently lack granular cell definitions (Figure 1B). In addition, cellular species compositions correlated between organs, ranging from low to high depending on the organs, and the simulated organ correlation distribution was similar to the Tabula Sapiens data (Figure 2D, Figure S1C).

Since there is no consensus model to estimate species richness (hereafter referred to as cellular richness), especially for cellular ecosystems, we first compared the accuracy of various non-parametric statistical estimators (see Methods). We assessed the performance of four estimators by drawing differently sized triplicate samples to assess the number of unique cellular species present in the ground truth simulated organ. We performed this for organs with distinct and partly overlapping cellular compositions. Notably, even in large organs with many rare cellular species, all the tested estimators started to stabilise towards the ground truth richness level after sampling 0.001% of the population, indicating that the estimators can efficiently estimate the true population value from a relatively small sample size, when 3 samples are available (Figure 2E). As expected, using more than 3 samples further enhanced the estimation accuracy (Figure S1D). Among the tested estimators, Chao2 showed the most promising results by being the fastest to plateau towards the true population value, while not being prone to overestimation. We also studied the error percentage for each cellular richness estimator by drawing differently sized samples from the simulated organs (Figure 2F). Notably, the Chao2 estimator in this case resulted in a median reduction of 18% error, compared to the estimates based solely on observed cellular species (species recorded in the samples). We therefore selected Chao2 for further use in case studies due to its desirable statistical properties.

As a conclusion, this simulation study and the benchmarking of existing statistical species richness estimators show that cellular richness estimation of organ systems is possible from existing, lognormally distributed single-cell atlas data, when 3 or more samples from a population are available. Importantly, the best-performing Chao2 estimator is simple to use and offers favourable statistical properties when estimating cellular richness from a lognormally distributed population.

### Estimating the completeness of large single-cell sequencing projects

Our previous results indicated that it is possible to confidently estimate the cellular richness using sample sizes of 0.001% or larger of the whole cellular population. Since many ongoing large-scale tissue profiling projects still fall below this percentage, we next wanted to assess in more detail the cellular richness of an organ, using repeated sampling from a large-scale single-cell data set. We argue that estimating the cellular richness of an organ and comparing it with the currently detected amount of cellular species allow for completeness estimation of any large-scale single-cell sequencing project.

To assess the completeness of a single-cell sequencing project, we used the single-cell data of neurotypical human brains provided by Emani et al. (Emani et al., 2024), which provides 17 good quality snRNA profiles, sampled from anterior, middle and posterior parts of the human dorsolateral prefrontal cortex (DLPFC), totalling 79,586 cells. Since we estimate the human prefrontal cortex having approximately 21 billion cells in total (Azevedo et al., 2009; McBride et al., 1999), the size of the Emani et al. dataset thus covers 0.0004% of the whole human prefrontal cortex. After filtering out doublets and poor quality cells using standard metrics (see Methods for details), and removing batch effects between the samples using the BBKNN method (Polański et al., 2020), the cells were clustered and displayed as a UMAP, showing marked differences in cellular complexity across the 17 samples (Figure 3A). Notably, even smaller cell clusters were often detected in multiple samples, indicating high completeness and good mixing between the different samples of the dataset (Figure 3A, labelled clusters). We evaluated the quality of batch effect removal by finding prefrontal cortex specific cell types based on RNA markers, confirming they are found from single clusters (Figure 3B).

**Fig. 3.**
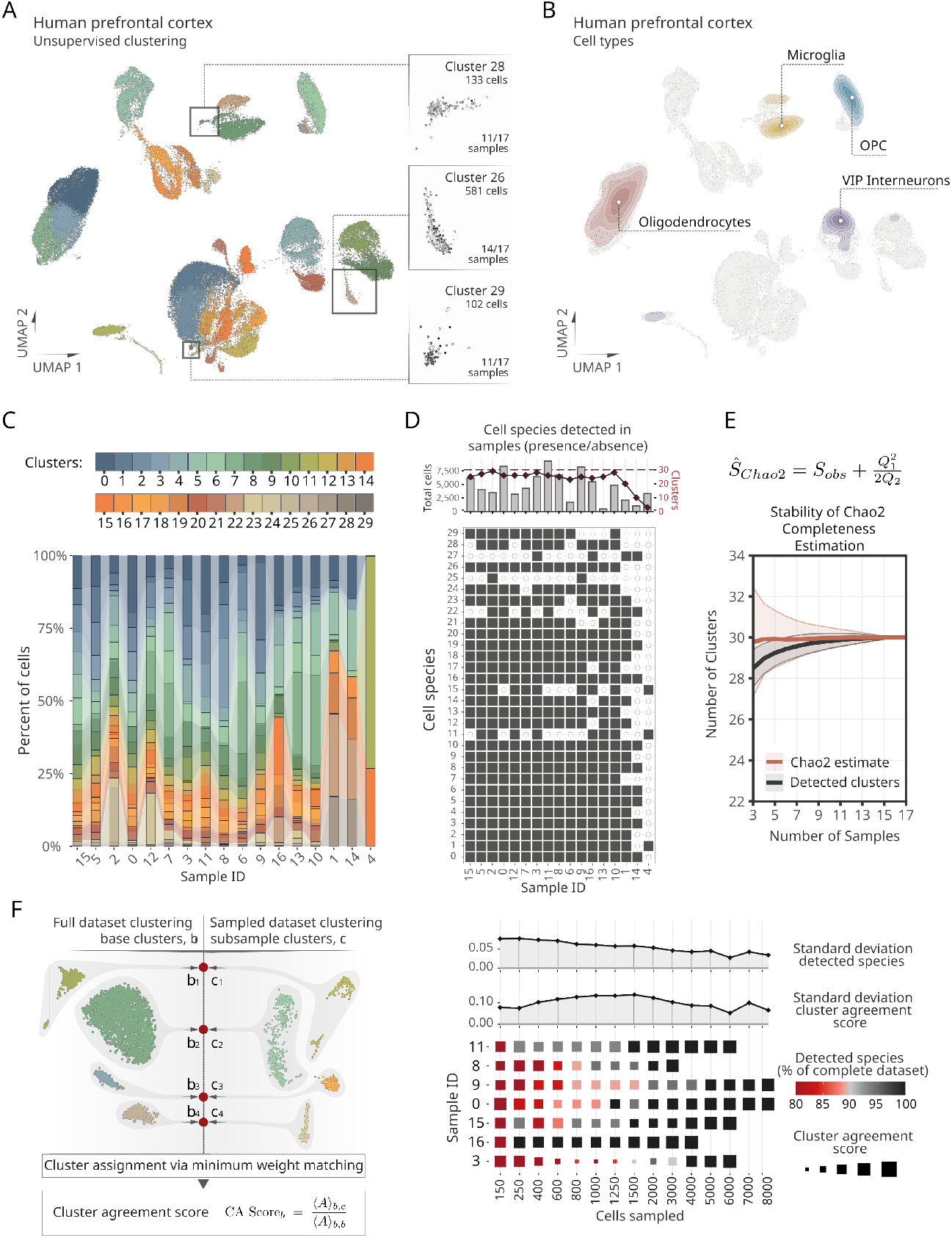
Completeness estimation of a large single-cell human prefrontal cortex dataset. (A) Dataset of single nuclei isolated from neurotypical human prefrontal cortices (17 samples in total from Emani et al., 2024). Colors indicate different Leiden clusters (30 distinct clusters detected across the 17 samples). Smaller clusters labelled within the integrated dataset were detected in multiple samples, indicating high dataset completeness. Right, inset panels (Clusters 28, 26 and 29; the three smallest detected clusters in terms of cell number); different grey tones correspond to the 17 individual samples. Transcriptionally similar cells from different samples converge within the same low-dimensional space, indicating shared biological cell states. (B) Cell type kernel density scoring of the human prefrontal cortex cell types. Module scores for four cell types commonly found in the human prefrontal cortex (oligodendrocytes, microglia, oligodendrocyte-progenitor cells (OPC), and VIP-expressing interneurons), estimated from their marker-gene set (see Methods), providing a density map that indicates the regions enriched for each cell type. (C) Cellular richness of each sample. Cellular species correspond here to the detected cell clusters. Samples are sorted by the standard deviation of their cluster proportions, starting with those showing the least variability. (D) Total number of cells and clusters detected in each sample (top) and the cell species presence/absence plot in each sample (bottom). (E) The full dataset was subsampled and the dynamics of cell cluster detection and Chao2 estimate was assessed using 100 random subsamples at each sample size. The traces correspond to the average number of clusters detected or estimated, and the confidence intervals to standard deviation. In the Chao2 Equation, *S*_*obs*_ is the observed cellular species count, *Q*_1_ the number of species observed in exactly one sample, and *Q*_2_ the number of species observed in exactly two samples. (F) Schematic workflow for cellular species detection stability. Standard procedure for single cell clustering (cell species detection) is applied to the full and subsampled datasets, yielding base clusters *b*_*i*_ and candidate clusters *c*_*j*_. Average *k*-NN connectivity mapping between every (*b*_*i*_, *c*_*j*_) pair is assembled into a cost matrix, and the Hungarian minimum-weight matching algorithm was used to select the single subsample cluster that best matches each base cluster. For each base cluster *b*, the cluster-agreement (CA) score compares the cluster connectivity of (*b, c*) to the within-cluster connectivity of (*b, b*) (see Equation), where ⟨*A*⟩_*u,v*_ is the mean *k*-NN edge connectivity between clusterings *u* and *v*. High CA scores indicate that the structure of base clustering is preserved in the subsample, whereas low scores indicate fragmentation or merging. (G) Stability of cellular species detection at different sample sizes. Seven samples with the highest single-cell coverage were selected for the analysis. Cluster stability analysis was performed by repeated subsampling and re-clustering. Upper panels display the standard deviation of detected species (%) and cluster agreement scores, respectively. The lower panel illustrates the average percentage of detected cellular species as a percentage of the complete dataset (color coding), and cluster agreement score (indicated by size) at different sample sizes.

Species richness estimation requires a criteria on how the cell types - here termed cellular species - are defined. Due to the absence of a broadly accepted definition of cell type, we define cellular species as transcriptionally different clusters found by unsupervised analysis of single-cell data. This criterion facilitates the systematic evaluation of cellular diversity across single-cell sequencing datasets. We first assessed the contribution of each sample on the completeness of the full dataset by analyzing the distribution and abundance of each cellular species within and across the samples (Figure 3C). This allowed us to assess both the consistency of cell species detection across samples and the extent of rare or sample-specific cell populations. As expected, the majority of cellular species were detected in most of the samples due to their larger size. However, we also observed sample-unique cellular clusters that were absent in most of the other samples (Figure 3C-D). Interestingly, our results showed that individual samples can already reveal much of the tissue’s complexity, trending towards the true species richness value (here 30 cell species) (Figure 3D).

To address the completeness of the prefrontal cortex dataset, we used the Chao2 estimator to estimate the number of cellular species in the human prefrontal cortex using all the 17 samples available. Although the coverage of this dataset was modest (0.0004%), applying the Chao2 estimator on the human prefrontal cortex showed that the estimates quickly stabilise into 30 cellular species (Figure 3E). This suggests that when the cellular species in the human prefrontal cortex are defined as here (i.e., all detected cell clusters), single-nucleus profiling of more samples will not reveal additional cellular species. This is further supported by the simulation studies showing how applying the Chao2 estimator on a dataset of 17 samples can estimate the complexity of even a highly complex organ with less than 1% error (Figure S1D). We therefore consider that the healthy human dorsolateral prefrontal cortex has been fully profiled. Importantly, we showed how Chao2 was able to correctly estimate the cellular richness of the full dataset, on average, already when applied to 3 or more samples (Figure 3E). However, increasing the number of samples led to more accurate estimations (narrower confidence intervals).

To further investigate how the sampling depth contributes to cellular richness estimation, we performed a more granular cluster stability analysis via subsampling and re-clustering of each sample (Figure 3F, see Methods). This approach allowed us to evaluate how robustly cellular species were detected at different sampling depths, identifying the point at which additional increases in analysed single cells provided diminishing returns in respect to new species detection. For the seven high-coverage samples, species recovery rates consistently exceeded 95%, and median cluster-agreement (CA) scores were greater than 0.95 when 3,000 - 4,000 cells per sample were analyzed. After reaching this threshold, further increasing the number of cells resulted in only minimal gains in CA scores and species recall rates, indicating that a practical saturation had been reached. Combining coverage-based stability estimates with Chao2 richness predictions thus offers a coherent strategy for guiding atlas-scale projects toward comprehensive and efficient characterization of cellular diversity.

In summary, the simulations and real data analyses together suggest that surprisingly low sample coverages can reveal all transcriptomically distinct cell species in a tissue, as long as multiple samples are available. Similar analysis can be applied to any ongoing single-cell sequencing or large-scale atlas projects,such as the Yao et al. mouse isocortex dataset (Yao et al., 2023), where 4 samples already captured the full transcriptional complexity (Figure S2A-E and Methods), hence providing estimates of their current completeness stage.

## Conclusions

Species richness estimation is a fundamental problem in ecology and biodiversity studies, where the goal is to determine the total number of species within a given habitat or ecosystem. The problem arises with the practical challenge of trying to observe all species, either plant or animal, which is in most cases difficult or impossible. The era of single-cell biology makes it relatively easy to understand tissues through their cellular composition, but similar to biodiversity studies, assessing the complexity of the whole tissue remains challenging. Previous works have estimated different aspects of the human cell populations, such as cellular size distribution and the number of total cells in the body (Bianconi et al., 2013; Hatton et al., 2023). In contrast, we focused here on leveraging single-cell datasets and statistical species richness estimation to provide a framework that enables cellular richness estimation (the number of different cell species in an organ). In particular, we provided methods for completeness estimation of any large-scale single-cell profiling project.

Currently, substantial investments are being made in mapping various cell types, their locations, and gene regulation across human tissues using single-cell RNA sequencing and related technologies. Although single-cell and single-nucleus RNA sequencing are currently the most practical and scalable technologies for defining cell types and their states, there exists versatile technologies and criteria on how cell types and cell states can be defined, and how to separate these two from each other (Dance, 2024). In addition, different protein-based techniques which do not suffer from RNA drop-outs are rapidly emerging (Unterauer et al., 2024). Currently, the cell type and state definitions are hard to disentangle using only data-driven approaches, especially in heterogeneous and dynamical tissue systems. Although it is likely that future advancements in technology and integrative multi-modal analysis will lead to sharper and more universally accepted definitions, today there still exist multiple definitions of cell types and states. Therefore, here we simply refer to all the detected cellular clusters as to cellular species. In future applications, it is possible to perform the same analytics and statistical estimations using more nuanced definitions of a cell type based on the emerging single-cell proteomics and multi-omics technologies.

Despite the significant interest in profiling various types of healthy and diseased tissues, there has so far been little efforts to estimate the total number of unique cellular species within these systems. Most studies have focused on cataloging cell populations based on the cells observed in a given dataset, without addressing the question of how many distinct cellular species—encompassing both cell types and cell states—remain undiscovered. Because a sample is almost always a poor estimate for the true cellular complexity of a tissue, we here sought to benchmark the available species richness estimators, their utility and sampling requirements to understand cellular diversity in different tissue systems. Although there exists extensive reviews about species richness estimators in other applications (Schmitz & Rahmann, 2024), herein we focused on finding estimators that would work especially with the single-cell profiling data. We identified Chao2 estimator as a simple and accurate method that offers favourable statistical properties when estimating cellular richness from a lognormally distributed population.

Our analyses suggest that the cellular species in most, if not all, human tissue systems are lognormally distributed. This led us to test Chao2 and other species richness estimators on lognormally distributed data, showing that many of the estimators are robust when the sample sizes are large enough. Importantly, such sample sizes are already available in many existing single-cell projects, particularly in large-scale initiatives, such as the Human Cell Atlas, and various tissue-specific profiling efforts, which often include millions of cells. To apply these statistical estimators on real-world data, we made use of a previously published, high-quality mouse and human datasets with multiple samples. These datasets revealed notable heterogeneity in cellular composition between the samples, strongly advising towards using multiple samples when attempting to assess cellular biodiversity. Generally, our analyses show that surprisingly shallow sample coverages can fully capture the cellular communities of an organ. Furthermore, performing many rounds of richness estimations when more samples become available is expected to make the estimates more accurate.

We reasoned that it is possible to understand single-cell atlas resources as a sample of the population and make statistical inference of the human body based on these somewhat large samples - similar to ecological populations. However, unlike for animal species, cells exhibit many favourable statistical properties. In particular, the probability of finding a cell from a tissue sample mainly depends on the abundance of the cell, which is especially true for single-nucleus profiling technologies. In addition, assessing sample coverage for cells is much easier than for animal species, which is expected to lead to more robust estimates. Importantly, the estimates of a dataset or atlas completeness are especially useful when building large single-cell resources, and we therefore suggest continuously running species richness estimates in regular intervals on every profiled organ, with the aim of understanding the level of completeness for each organ that is being profiled. As single-cell resources continue to expand and the profiling depths across organs become more balanced, this work will provide the basis of future estimates of even the whole human body.

## Supplementary Note 1: Methods

### A. Simulations of organs and human

Since the organs are partially correlated, we simulated a human body with a defined number of organs (*n*=20), and cell types by drawing a sample of size 5 trillion cells from a multivariate log-normal distribution. To define organ variable dependencies, the covariance structure of this distribution was derived from the empirical covariance matrix of Tabula Sapiens. The user-defined parameters in the simulation were:

- **n**: number of cells in the whole organism
- **m**: number of organs
- **k**: number of cell types

To simulate a human body with an organ system, we first generated data in log-transformed space, where the organ abundances followed a multivariate normal distribution. This was done using the mvrnorm() function from the MASS library in R. We then applied an exponential transformation to obtain log-normally distributed organ abundances.

To simulate realistic dependencies between organs, we analyzed Tabula Sapiens data, assessing its organ means and co-variance structure in both normal and log-normal spaces. We found that the means of log-transformed organ abundances followed a normal distribution, allowing us to simulate a corresponding vector of means for the normally-distributed space. However, because exponentiation is a nonlinear transformation, it alters dependencies between the variables.

To retain the empirical covariance structure after back-transformation, we derived a mathematical relationship between the covariance matrices in normal and log-normal spaces. For this, we considered Y as a multivariate log-normal random vector:

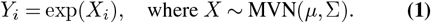

The covariance for two dependent variables (*Y*_*i*_, *Y*_*j*_) in logarithmic space is by definition:

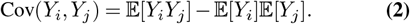

Because *Y*_*i*_ = exp(*X*_*i*_), the expected value of *Y*_*i*_*Y*_*j*_ is:

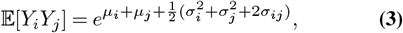

where *σ*_*ij*_ is the covariance in normal space. The relationship between and the covariance in log-normal space depends on the covariance in normal space:

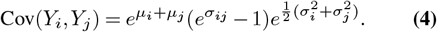

Rearranging equation (4), we can express covariance in normal space as:

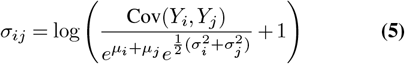

This enables covariance matrix (Σ) back-transformation while retaining dependencies between the variables.

### B. Species Richness Estimators

We reasoned that statistical estimators which use presence/absence data, rather than abundance data, are more suitable for the use with single-cell or single-nucleus RNA sequencing data. Many abundance data-based estimators rely on singlet-detection, which can be difficult with current RNA-based methods. Instead, we focused on benchmarking statistical estimators which use presence/absence data and that rely on repetitive sampling.

### Chao2 (For Incidence Data)

The Chao2 estimator is a non-parametric method for estimating species richness based on incidence data. It accounts for unseen species by leveraging the frequency of rare species across samples.

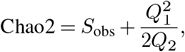

where *S*_obs_ is the observed species count, *Q*_1_ is the number of species observed in exactly one sample, and *Q*_2_ is the number of species observed in exactly two samples (Chao, 1984).

### ICE (Incidence-based Coverage Estimator)

The ICE estimator is a non-parametric statistical method that uses presence/absence data to infer the number of unseen species. It emphasizes the contribution of rare species across samples (Chao, 1987).

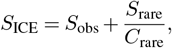

where:

- *S*_ICE_: the estimated total species richness.
- *S*_obs_: the number of species observed in the dataset.
- *S*_rare_: the number of species classified as rare (by default those appearing in ≤ 10 samples).
- *C*_rare_: the sample coverage estimate for rare species, calculated as:

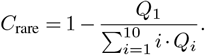

Here:

- *Q*_1_: the number of species that occur in only one sample.
- *Q*_*i*_: the number of species that occur in exactly *i* samples.

### Jackknife1 (For Incidence Data)

The Jackknife1 estimator is a resampling technique for estimating species richness. For incidence data, it is calculated as:

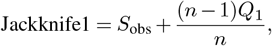

where *S*_obs_ is the observed species count, *n* is the number of samples, and *Q*_1_ is the number of species observed in exactly one sample (Heltshe & Forrester, 1983).

### Jackknife2 (For Incidence Data)

The Jackknife2 estimator refines the richness estimate by considering species observed in only one or two samples. It is calculated as:

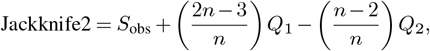

where *Q*_1_ and *Q*_2_ are the number of species observed in exactly one and exactly two samples, respectively (Heltshe & Forrester, 1983).

### C. Statistical Methods

Goodness of fit testing for lognormal distributions was done using the Kolmogorov-Smirnov test. All the statistical analyses as well as simulations were run using `R` with `R Studio`. Different statistical estimators were used through `R` with the CRAN package `fossil` (Vavrek, 2011).

### D. Single-Cell Analytics

The single-cell analysis was performed using **Python** and **Scanpy** (Wolf *et al*., 2018). In brief, raw sequencing data were filtered to retain only cells with mitochondrial gene content below 5% and at least 200 detected genes. Human prefrontal cortex samples, which exhibited higher transcriptional heterogeneity, were integrated using **scVI** (Gayoso *et al*., 2022), and cell doublets were identified and removed with **Scrublet** (Wolock *et al*., 2019). One sample from the human prefrontal cortex dataset was removed in the beginning for having less than 10 cells. Clustering of the data was done using Scanpy’s Louvain algorithm. For assessing data integration between samples, the cell type scores were calculated by averaging the expression of a set of RNA markers for each cell type.

### Marker Genes for Human Prefrontal Cortex

- **Oligodendrocytes**: MOG, MBP, CNP, PLP1, MAG
- **Microglia**: PTPRC, CSF1R, AIF1, CX3CR1, P2RY12
- **Oligodendrocyte Progenitor Cells (OPCs)**: PDGFRA, CSPG4, OLIG1, OLIG2, SOX10
- **VIP Interneurons**: VIP, CALB2, HTR3A, RELN, TAC3

### Marker Genes for Mouse Isocortex

- **Inhibitory Neurons (GABAergic neurons)**: Gad1, Gad2, Pvalb, Sst, Vip, Nkx2-1
- **Oligodendrocytes**: Mog, Mbp, Cnp, Plp1, Olig1
- **Endothelial Cells**: Pecam1, Cldn5, Kdr

Module scores were computed using Seurat’s `AddModuleScore` (Hao *et al*., 2023). After scoring, we estimated a weighted kernel-density surface of cells on the UMAP embedding, using each cell’s module score as the weight. The background density was then subtracted, yielding a density map that indicates regions enriched for each cell type.

### E. Data Integration and Batch Effect Correction for Mouse Brain Data

To integrate single-cell data and correct for batch effects in the mouse isocortex data, we trained a **single-cell Variational Inference (scVI)** model (Lopez *et al*., 2018) on the human prefrontal cortex dataset with the following parameters:

- **Latent space dimension** (n_latent): 30
- **Hidden layer sizes** (n_hidden): 128
- **Number of hidden layers** (n_layers): 2
- **Dropout rate** (dropout_rate): 0.1
- **Likelihood model** (gene_likelihood): “zinb” (default in scVI)
- **Batch correction** (use_batch_norm, use_layer_norm): Enabled via key=‘sample’
- **Library size scaling** (use_observed_lib_size): True
- **Learning rate** (lr): 1 *×* 10^−3^
- **Number of epochs** (max_epochs): 500
- **Batch size** (batch_size): 256
- **KL-weight annealing**: Default scVI schedule
- **Early stopping**: Patience = 20
- **Dispersion modeling** (dispersion): “gene-batch”

### F. Data Integration and Batch Effect Correction for Human Prefrontal Cortex

To integrate single-cell data and correct for batch effects in the human prefrontal cortex dataset, we used **Batch-Balanced K-Nearest Neighbors (BBKNN)** (Polański *et al*., 2020) with these parameters:

- **Neighbors within batch**: 7
- **Number of principal components**: 25
- **Trim parameter**: 30

### G. Cluster Stability Analysis

The cluster stability analysis was performed via subsampling and re-clustering of each sample. Each sample was first clustered (defining “base clusters”), after which multiple subsamples of varying sizes were independently re-clustered. For each subsample:

1. PCA was recomputed.
2. A *k*-nearest neighbors (k-NN) connectivity graph was constructed with *k* set proportional to subsample size (minimum *k* = 5).
3. Reclustering was performed using the Louvain algorithm.

Subsample clusters were matched to base clusters via the Hungarian minimum-weight assignment algorithm (from `scipy.optimize`), maximizing average k-NN connectivity. Cluster agreement (CA) scores quantified stability as

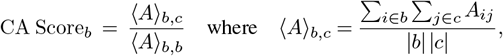

where, *A*_*ij*_ denotes k-NN edge connectivity between cells *i* and *j*, and *b* and *c* represent matched clusters from the base and subsampled data, respectively. The ratio of subsample-to-base cluster similarity was then aggregated across subsample sizes and visualized as a stability heatmap.

### H. Data Availability

The codes to produce the simulated data are available on GitHub (https://github.com/mitroe/AtlasCompleteness). The real-world cell type and single-cell data were extracted from the following open databases:

- **[Tabula Sapiens](https://tabula-sapiens.sf.czbiohub.org/)**: data accessed on 17.05.2024
- **[CELLxGENE](https://cellxgene.cziscience.com/)**: data accessed on 30.07.2024
- **Human cell treemap, Organ size estimates**: data accessed on 30.05.2024
- **Human prefrontal cortex (Emani *et al***., **2024)**: raw data available on Gene Expression Omnibus with the accession number GSE261983
- **Mouse isocortex (Yao *et al***., **2023)**: raw and processed data available at https://assets.nemoarchive.org/dat-qg7n1b0

### I. Code Availability

All code used to produce the analyses shown in this publication are available on [GitHub](https://github.com/mitroe/AtlasCompleteness/tree/main/Supplements).

### J. Author Contributions

Conceptualisation: T.A., M.M. Data curation: S.V. Formal analysis: M.M., Y.C., S.V., Y.A. Funding acquisition: T.A., M.M. Investigation: T.A., M.M., Y.C., S.V., Y.A. Methodology: T.A., M.M., Y.C., Y.A. Project administration: M.M. Resources: T.A. Supervision: T.A., M.M. Validation: M.M., Y.A., Y.C. Visualization: M.M., Y.A. Writing – original draft: T.A., M.M., Y.A. Writing – review & editing: T.A., M.M.

## Supplementary material

Supplementary material can be found from here, or at the end of this document.

## Declaration of Interests

The authors declare no competing interests.

## Acknowledgements

We thank Henri Pesonen, Leo Lahti, and Yunus Vilkkavaara-Cankocak for expert comments during the writing of the manuscript. All single-cell data analytics were performed using a computing cluster. The authors acknowledge the CSC – IT Center for Science, Finland, for providing the computational resources. MM was supported by the Sakari Alhopuro Foundation. TA was supported by the Norwegian Cancer Society (grants 216104 and 273810), Norwegian Health Authority South-East (grants 2020026 and 2023105), the Radium Hospital Foundation, the Finnish Cancer Foundation, Sigrid Jusélius Foundation, the Research Council of Finland under the frame of EP PerMed (CLL-OUTCOME, grant 367855), and the Research Council of Norway under the frame of EP PerMed (ImmuneT-ME, grant 357095).

## SUPPLEMENTARY INFORMATION

**Supplementary Figure 1.**
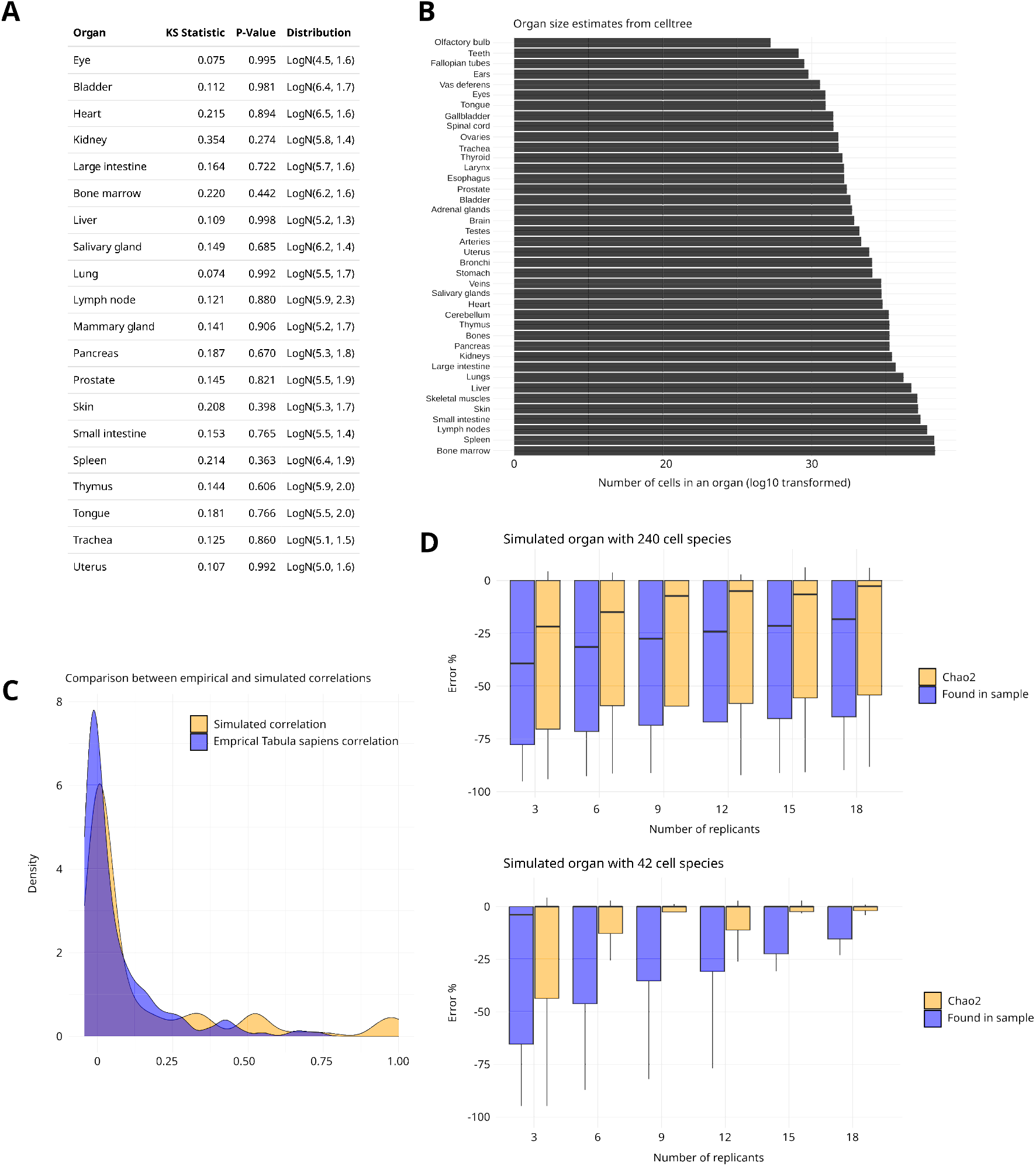
(A) Goodness-of-fit testing whether cell type abundances from different organs are log-normally distributed. The table reports test statistics from Kolmogorov-Smirnov (KS) test and the corresponding p-values. Note: high p-values indicate insufficient evidence to reject the null hypothesis of log-normal distribution. (B) The estimates of total cell counts in each organ (from https://humancelltreemap.mis.mpg.de/). (C) Comparison of empirical correlations between Tabula Sapiens data and simulated human organs. KS test for distribution similarity, with p = 0.42, indicating similar distributions. (D) Effect of the number of sample replicates on the Chao2 estimation accuracy. The horizontal lines indicate median error levels from -100% (total underestimation) to 0% (no error), boxes the interquartile ranges, and the vertical lines indicate the number of samples.

**Supplementary Figure 2.**
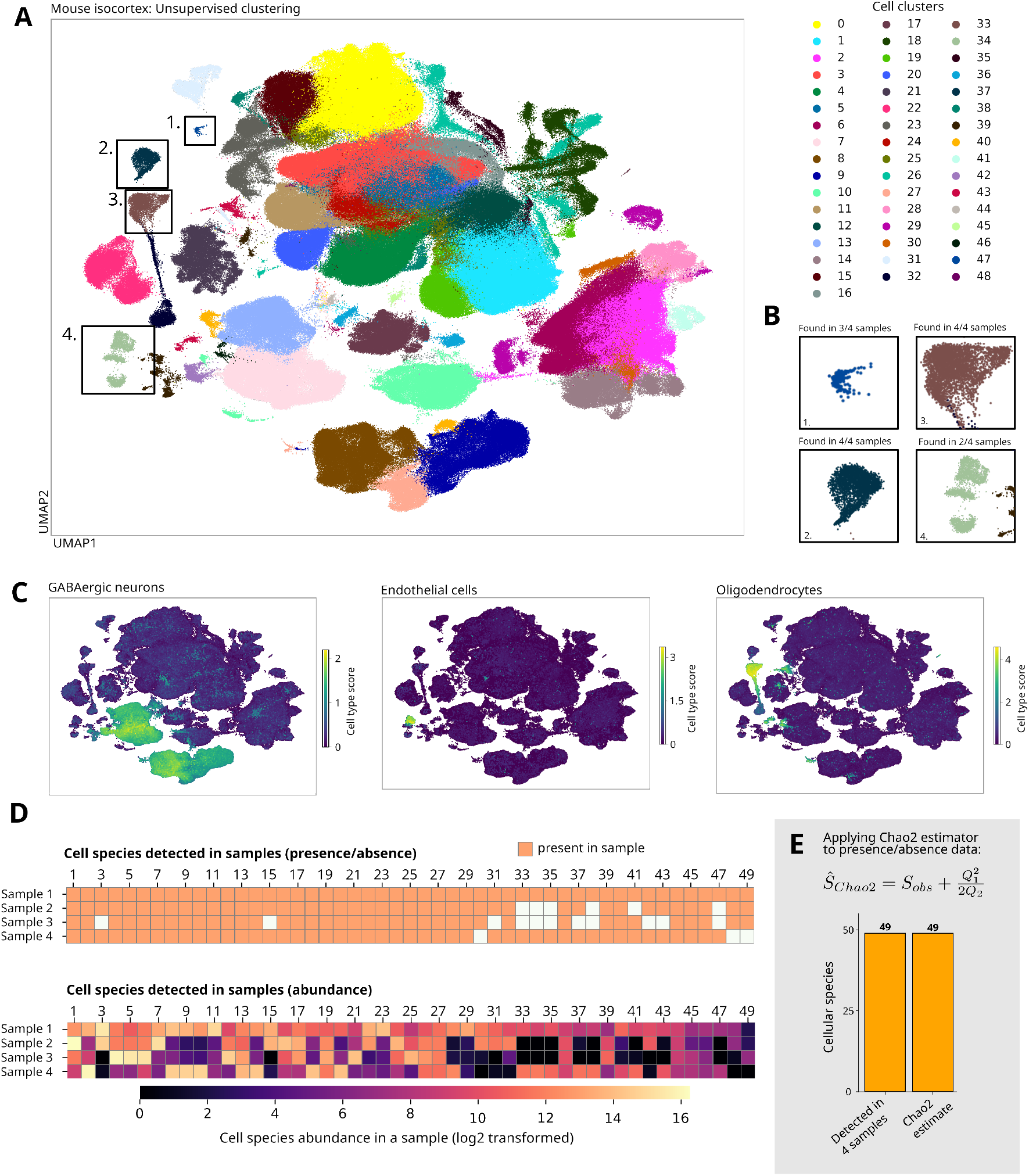
Completeness estimation of a large single-cell mouse isocortex dataset. (A) one million cell dataset with 4 individual samples analyzed with 10x chromium V2 chemistry of the mouse isocortex from Yao et al., 2023. Leiden clusters are shown. Colors indicate different clusters (49 distinct clusters detected across 4 samples). (B) Smaller clusters labelled in panel A within the integrated dataset are detected in multiple samples indicating high dataset completeness. (C) Individual clusters identified with RNA markers and matched to common cell types found in mouse isocortex. The cell type score is calculated by averaging gene expression values of the marker genes separately for each cell type (see Methods). (D) Cellular species detection in the four isocortex samples (upper panel: presence/absence and lower panel: abundance data). Cellular species correspond here to the detected cell clusters. (E) Chao2 estimate of the cellular richness of mouse isocortex dataset using data from Yao et al. In the Chao2 Equation, *S*_*obs*_ is the observed cellular species count, *Q*_1_ the number of species observed in exactly one sample, and *Q*_2_ the number of species observed in exactly two samples.

## Notes

https://github.com/mitroe/AtlasCompleteness

### Competing Interest Statement

The authors have declared no competing interest.

### Summary of Updates

Completely new figures and new results for figures 1 and 3. Also the text is vastly improved

## Bibliography

1. Azevedo, F. A. C., Carvalho, L. R. B., Grinberg, L. T., Farfel, J. M., Ferretti, R. E. L., Leite, R. E. P., Jacob Filho, W., Lent, R., & Herculano-Houzel, S. (2009). Equal numbers of neuronal and nonneuronal cells make the human brain an isometrically scaled-up primate brain. The Journal of Comparative Neurology, 513(5), 532–541. 10.1002/cne.21974

2. Bianconi, E., Piovesan, A., Facchin, F., Beraudi, A., Casadei, R., Frabetti, F., Vitale, L., Pelleri, M. C., Tassani, S., Piva, F., Perez-Amodio, S., Strippoli, P., & Canaider, S. (2013). An estimation of the number of cells in the human body. Annals of Human Biology. https://www.tandfonline.com/doi/abs/10.3109/03014460.2013.807878

3. Chao, A. (1984). Nonparametric Estimation of the Number of Classes in a Population. Scandinavian Journal of Statistics, 11(4), 265–270.

4. Chao, A. (1987). Estimating the Population Size for Capture-Recapture Data with Unequal Catchability. Biometrics, 43(4), 783–791. 10.2307/2531532

5. Chiu, C.-H. (2023). Sample coverage estimation, rarefaction, and extrapolation based on sample-based abundance data. Ecology, 104(8), e4099. 10.1002/ecy.4099

6. Dance, A. (2024). What is a cell type, really? The quest to categorize life’s myriad forms. Nature, 633(8031), 754–756. 10.1038/d41586-024-03073-2

7. Emani, P. S., Liu, J. J., Clarke, D., Jensen, M., Warrell, J., Gupta, C., Meng, R., Lee, C. Y., Xu, S., Dursun, C., Lou, S., Chen, Y., Chu, Z., Galeev, T., Hwang, A., Li, Y., Ni, P., Zhou, X., Bakken, T. E., … Gerstein, M. (2024). Single-cell genomics and regulatory networks for 388 human brains. Science, 384(6698), eadi5199. 10.1126/science.adi5199

8. Gayoso, A., Lopez, R., Xing, G., Boyeau, P., Valiollah Pour Amiri, V., Hong, J., Wu, K., Jayasuriya, M., Mehlman, E., Langevin, M., Liu, Y., Samaran, J., Misrachi, G., Nazaret, A., Clivio, O., Xu, C., Ashuach, T., Gabitto, M., Lotfollahi, M., … Yosef, N. (2022). A Python library for probabilistic analysis of single-cell omics data. Nature Biotechnology, 40(2), 163–166. 10.1038/s41587-021-01206-w

9. Grilli, J. (2020). Macroecological laws describe variation and diversity in microbial communities. Nature Communications, 11(1), 4743. 10.1038/s41467-020-18529-y

10. Gwinn, D. C., Allen, M. S., Bonvechio, K. I., Hoyer, M. V., & Beesley, L. S. (2016). Evaluating estimators of species richness: The importance of considering statistical error rates. Methods in Ecology and Evolution, 7(3), 294–302. 10.1111/2041-210X.12462

11. Hao, Y., Stuart, T., Kowalski, M. H., Choudhary, S., Hoffman, P., Hartman, A., Srivastava, A., Molla, G., Madad, S., Fernandez-Granda, C., & Satija, R. (2023). Dictionary learning for integrative, multimodal and scalable single-cell analysis. Nature Biotechnology, 1–12. 10.1038/s41587-023-01767-y

12. Hatton, I. A., Galbraith, E. D., Merleau, N. S. C., Miettinen, T. P., Smith, B. M., & Shander, J. (2023). The human cell count and size distribution. Proceedings of the National Academy of Sciences, 120(39), e2303077120. 10.1073/pnas.2303077120

13. Heltshe, J. F., & Forrester, N. E. (1983). Estimating Species Richness Using the Jackknife Procedure. Biometrics, 39(1), 1–11. 10.2307/2530802

14. Lopez, R., Regier, J., Cole, M. B., Jordan, M. I., & Yosef, N. (2018). Deep generative modeling for single-cell transcriptomics. Nature Methods, 15(12), 1053–1058. 10.1038/s41592-018-0229-2

15. Magurran, A. E. (1988). Ecological Diversity and Its Measurement. Springer Netherlands. 10.1007/978-94-015-7358-0

16. McBride, T., Arnold, S. E., & Gur, R. C. (1999). A comparative volumetric analysis of the prefrontal cortex in human and baboon MRI. Brain, Behavior and Evolution, 54(3), 159–166. 10.1159/000006620

17. Vavrek, M. J. (2011). fossil: palaeoecological and palaeogeographical analysis tools. Palaeontologia Electronica, 14(1), 1 T. R package version 0.4.0. Retrieved from http://palaeo-electronica.org/2011_1/238/238.pdf

18. Polański, K., Young, M. D., Miao, Z., Meyer, K. B., Teichmann, S. A., & Park, J.-E. (2020). BBKNN: Fast batch alignment of single cell transcriptomes. Bioinformatics, 36(3), 964–965. 10.1093/bioinformatics/btz625

19. Program, C. S.-C. B., Abdulla, S., Aevermann, B., Assis, P., Badajoz, S., Bell, S. M., Bezzi, E., Cakir, B., Chaffer, J., Chambers, S., Cherry, J. M., Chi, T., Chien, J., Dorman, L., Garcia-Nieto, P., Gloria, N., Hastie, M., Hegeman, D., Hilton, J., … Carr, A. (2023). CZ CELL×GENE Discover: A single-cell data platform for scalable exploration, analysis and modeling of aggregated data (p. 2023.10.30.563174). bioRxiv. 10.1101/2023.10.30.563174

20. Rood, J. E., Wynne, S., Robson, L., Hupalowska, A., Randell, J., Teichmann, S. A., & Regev, (2024). The Human Cell Atlas from a cell census to a unified foundation model. Nature, 1–2. 10.1038/s41586-024-08338-4

21. Schmitz, J. E., & Rahmann, S. (2024). A Review and Evaluation of Species Richness Estimation (p. 2024.10.09.615408). bioRxiv. 10.1101/2024.10.09.615408

22. Tabula Sapiens Consortium*, Jones, R. C., Karkanias, J., Krasnow, M. A., Pisco, A. O., Quake, S. R., Salzman, J., Yosef, N., Bulthaup, B., Brown, P., Harper, W., Hemenez, M., Ponnusamy, R., Salehi, A., Sanagavarapu, B. A., Spallino, E., Aaron, K. A., Concepcion, W., Gardner, J. M., … Wyss-Coray, T. (2022). The Tabula Sapiens: A multiple-organ, single-cell transcriptomic atlas of humans. Science, 376(6594), eabl4896. 10.1126/science.abl4896

23. Unterauer, E. M., Boushehri, S. S., Jevdokimenko, K., Masullo, L. A., Ganji, M., Sograte-Idrissi, S., Kowalewski, R., Strauss, S., Reinhardt, S. C. M., Perovic, A., Marr, C., Opazo, F., Fornasiero, E. F., & Jungmann, R. (2024). Spatial proteomics in neurons at single-protein resolution. Cell, 187(7), 1785–1800.e16. 10.1016/j.cell.2024.02.045

24. Wolf, F. A., Angerer, P., & Theis, F. J. (2018). SCANPY: Large-scale single-cell gene expression data analysis. Genome Biology, 19(1), 15. 10.1186/s13059-017-1382-0

25. Wolock, S. L., Lopez, R., & Klein, A. M. (2019). Scrublet: Computational Identification of Cell Doublets in Single-Cell Transcriptomic Data. Cell Systems, 8(4), 281–291.e9. 10.1016/j.cels.2018.11.005

26. Yao, Z., van Velthoven, C. T. J., Kunst, M., Zhang, M., McMillen, D., Lee, C., Jung, W., Goldy, J., Abdelhak, A., Aitken, M., Baker, K., Baker, P., Barkan, E., Bertagnolli, D., Bhandiwad, A., Bielstein, C., Bishwakarma, P., Campos, J., Carey, D., … Zeng, H. (2023). A high-resolution transcriptomic and spatial atlas of cell types in the whole mouse brain. Nature, 624(7991), 317–332. 10.1038/s41586-023-06812-z

27. Zeng, H. (2022). What is a cell type and how to define it? Cell, 185(15), 2739–2755. 10.1016/j.cell.2022.06.031

